# Novel pipeline of high-frequency neoantigens heathy donor-based validation in breast cancer

**DOI:** 10.1101/596908

**Authors:** Lili Qin, Ying Huang, Zhaoduan Liang, Geng Liu, Xiumei Lin, Ting An, Dongli Li, Le Cheng, Bo Li

**Affiliations:** College of Basic Medicine, Dali University, Dali 671000, China; BGI-Shenzhen, Shenzhen 518083, China; BGI-Yunnan, BGI-Shenzhen, Kunming, 650106, China; BGI-GenoImmune, BGI-Shenzhen, Wuhan, 430079, China; China National GeneBank, BGI-Shenzhen, Shenzhen 518120, China

**Keywords:** breast cancer, neoantigens, database, human healthy donor, immunogenicity

## Abstract

Neoantigen, a peptide fragment formed by genetic mutation, gives immunologist a new target for cancer therapy. Development of biotechnology has opened a new era for discovering high-frequency neoantigens. The aim of our study was to identify breast cancer neoantigens for tumor immunotherapy using an efficient way. Here, we established a computational pipeline to identify neoantigens associated with breast cancer using data from database and evaluated the immunogenicity of neoantigens using the peripheral blood of healthy donators in vitro. We identified 39,401 missense mutation sites from 285,283 single nucleotide variations (SNVs) obtained from database, and confirmed candidate epitopes by analyzing the binding affinity of mutant epitopes and human leukocyte antigen (HLA) using 6 algorithms. Peptide-binding assay was used as a complement for affinity testing. The immunogenicity of candidate peptides with high affinity were assessed through enzyme-linked immunospot (ELISPOT) assay and Cytotoxicity assay. In our study, we identified 10 candidate peptides with high binding affinity of HLA-A*0201 alleles, and seven of ten peptides showed the ability of inducing specific cytotoxic lymphocytes(CTLs) ex vivo, in healthy HLA-A2^+^ donors. We found that the peptide derived from TWISTNB have the highest immunogenicity and cytotoxicity among those candidate peptides. Furthermore, it can trigger the immune response of specific-CTLs to destroy target cells expressing this neoantigen in vitro, and without cross-reactivity with wild-type peptides. We conclude that the effective pipeline will provide potential possibilities to rapidly identify abundant high-frequency neoantigens and create neoantigen library for immunotherapy of breast cancer and even other tumors.

## 1. Introduction

Breast cancer is the most common tumor in female and there are more than 500,000 women worldwide died of this in 2011(Pham *et al.*, 2015). In 2018, a report on the cancer worldwide shown that incidence and mortality of breast cancer have been at the top among common cancers of females, 24.2% and 15% respectively (Bray *et al.*, 2018). Breast cancer is a highly heterogeneous disease caused by the accumulation of gene mutations, epigenetic disorders and other factors. In China, most cases of breast cancer are found to be advanced, therefore traditional surgery, radiotherapy and chemotherapy methods cannot achieve the desired therapeutic effect. In recent years, although molecular targeted drugs, endocrine therapy have obvious effect on the treatment of early breast cancer, the control of malignant progression of breast cancer is still a severe problem in the field of breast cancer treatment. Hence, it is urgent to find new molecular targets and develop new treatment methods.

Tumor immunotherapy resists and eliminates tumors by improving autoimmunity. At present, there are two effective immunotherapy methods. One is immune checkpoint block drugs, which can relieve the inhibitory pathway in the tumor microenvironment and trigger the immune system to recognize and destroy tumors. The development of PD1/ PDL1 is the fastest in the study of these immune checkpoints, the first clinical trial of anti-PD1/PDL1 agents began in 2006, reached 2,250 in 2018, and an increasing number of anti-PD1/PDL1 agents approved by the FDA for the treatment of various cancers(Tang *et al.*, 2018). The other one is adoptive immunotherapy, which adopted reinfusion of immune cells and made returned immune cells to recognize and eliminate tumors. In the past few years, following PD1/PDL1 therapy achieved striking efficacy, CAR-T (Chimeric Antigen Receptor T-Cell) immunotherapy has created a new miracle in anti-tumor immunotherapy. According to relevant clinical data: anti-CD19 CAR T cells were used to treat relapsed or refractory B cell malignancies achieve the CR(complete remission) in various trials range from 70% to 94%(Miliotou & Papadopoulou, 2018). Thence, a growing number of scholars commit to tumor immunotherapy and look forward to finding a broad and effective approach to cope with cancer. From 2006 to 2014, the number of clinical trials registered on ClinicalTrials.gov increased more than 9000, according to Association of Community Cancer Centers (ACCC)(https://www.accc-cancer.org/home/learn/immunotherapy/resource-detail/Clinical-Trials-in-Immunotherapy).

Neoantigens are generated from tumor-specific proteins encoded by somatic mutations and entirely absent from normal human tissue. They are presented by major histocompatibility complexes (MHCs) on surfaces of tumor cells or antigen presenting cells, recognized by T cells and induced specific CTLs to destruct the tumor but not the normal tissue(Tubb *et al.*, 2018). Neoantigens have a significant momentum in recent years, they are beneficial to assess the therapeutic effect of immunotherapy and offer targets for tumor immunotherapy (Wu *et al.*, 2018). Although neoantigens can produce a robust antigen-specific response, finding out and validating the high-frequency or shared neoantigens in various cancers still face enormous challenges. The rapid development of the next-generation sequencing technologies and bioinformatics has taken an essential role in neoantigens ascertainment. Hartmaier *et al*. identified shared neoantigens across cancers from 63,220 tumor genome sequencing data, revealing that there are some neoantigens exist in various cancers, such as *KRAS* G12C, *PIK3CA, EGFR* L858R (Hartmaier *et al.*, 2017). The establishment of a database for cancer genomics has facilitated our search for candidate neoantigens, such as TRON Cell Line Portal (TCLP), The Cancer Immunome Atlas (TCIA), International cancer genome consortium (ICGC) and The Cancer Genome Atlas (TCGA). Brown *et al*. used genomic data from the TCGA to characterize neoantigens and analysis the relationship between immunogenic neoantigens and patient survival, revealing that tumor bearing immunogenic mutations have elevated the expression of CD8A as well as CTLA-4 and PDCD1(Brown *et al.*, 2014). Methods of identifying tumor neoantigens in vitro using healthy donators for developing targets of TCR-T drugs have been reported. These candidate neoantigens were obtained from the non-synonymous mutation in sequencing data for clinical samples of tumors, verified in vitro using healthy donators-derived immune cells, and used as a target for TCR-T drug research(Kato *et al.*, 2018) or (Matsuda *et al.*, 2018).

This strategy identified high frequency candidate neoantigens in breast cancer from existing data in databases and used human healthy donor-derived CD8+T cells to verify neoantigens in vitro. Moreover, it provides a research basis for clinical application of tumor immunotherapy such as DC vaccine, peptide vaccine and DC-CTLs.

## 2. Materials and methods

### (i) Ethics statement

All donors were given written informed consent to participate this experiment. This study was approved by the Institutional Ethics Committee of BGI (Shenzhen, China).

### (ii) Cell lines and peptides

MHC class I-restricted peptides were synthesized at > 95 % purity at GenScript Biotech Corp (Nanjing, China) as confirmed by mass spectrometry. Human transporter associated with antigen processing (TAP)-deficient cell line (T2) was HLA-A*0201 positive and purchased from ATCC (CRL-1992, Rockville, USA), and cultured in IMDM (Gibco, USA) containing 10% fetal bovine serum (Hyclone, USA). HCT116 cell line (HLA-A*0201, colon carcinoma) were transferred from Guang Dong HEC Pharm Co Ltd, genetically modified-HCT116 was prepared by transfection with minigenes containing TWISTNB-derived peptides (KLMGIVYKV), and both of them were cultured in RPMI Medium 1640 (Gibco, USA) containing 10% fetal bovine serum.

### (iii) Neoantigens prediction

In this study, a prediction pipeline was established to find mutated epitopes in breast tumor samples from TCGA, ICGC and an article (https://www.nature.com/articles/nature11143). For the sake of maximizing the prediction accuracy, the capability for missense mutations of breast cancer binding to MHC was predicted using multiple algorithms: PSSMHCpan v1.0(Liu *et al.*, 2017), NetMHC v3.4(Sun *et al.*, 2014), NetMHCcons v1.1 (Karosiene *et al.*, 2012), NetMHCpan v2.8(Hoof *et al.*, 2009), Stabilized Matrix Method (SMM) (Peters & Sette, 2005) and the SMM with a Peptide:MHC Binding Energy Covariance algorithm (SMMPMBEC) (Kim *et al.*, 2009). Firstly, we annotated missense variants in SNVs with ANNOVAR to tick off candidate epitopes (length was 9 mer) with an in-house script. Then, we predicted the affinity of HLA-A*0201 mutant epitopes to MHC class I using above 6 algorithms. Results were used as binding affinity, IC50<500nM(half-maximum inhibitory concentrations) at least in two algorithms(Johanns *et al.*, 2016). The lower of the IC50, the stronger of the affinity. Finally, the IC50 is standardized as the binding score. The higher of the score, the stronger of the affinity.

### (iv) Peptide-binding assay for neoantigens

We used T2 cells to confirm the binding affinity of candidate peptides and HLA. Following a previously protocol (Hansen & Myers, 2003) with some modifications. 2 ×10^5^ T2 cells were cultured in IMDM medium without serum, peptides were added to T2 cells at the final concentration 100ug/ml, and then cultured at 37°C in a CO_2_ incubator overnight (16h-20h). In our methods, the Melan-A_26-35_(ELAGIGILTV) was used as the positive peptide, and without peptide as the negative control. Subsequently, cells were stained by anti-human HLA-A2 antibody (BB7.2, BioLegend, USA) with PE-labeled. FI (Fluorescence index) value was used as the evaluation standard. Refer to previous assessment criteria, FI = (mean FITC fluorescence for the given peptide-mean FITC fluorescence without peptide) / (mean FITC fluorescence without peptide) (Lv *et al.*, 2009).

### (v) Induction of neoantigens specific CTLs in vitro from healthy donor

To induce neoantigens specific CTLs, we first isolated the peripheral blood mononuclear cells (PBMCs) from healthy donors by Ficoll-Paque PLUS density gradient centrifugation (GE Healthcare, USA), CD8^+^ T cells were sorted from PBMC by CD8 MicroBeads (human) (Miltenyi, Germany) and were cryopreserved until use. Then the PBMC without CD8^+^ T cells were used to generate monocyte-derived dendritic cells (DCs) by plastic adherence culture method. DC precursor cells were seeded into plates in AIM-V medium (Gibco, USA) supplemented with 5% autologous serum, 80ng/ml GM-CSF (Gibco, USA), 100ng/ml IL-4 (Gibco, USA) for 48h, and then added half amount of medium contain 1ug/ml CD40L(Peprotech, USA) and 250u/ml IFN-γ(Peprotech, USA)to induce monocyte differentiated into DCs and matured for 48h. DCs were pulsed with 10ug/ml candidate peptides overnight and recovered cryopreserved CD8^+^T cells rest for overnight. Following the day, DCs were collected and cocultured with autologous CD8^+^ T cells with the ratio between 1:4 to 1:10 in AIM-V medium with 5% autologous serum and 30 ng/ml IL-21 (Cellgenix, Germany). Two days later, human recombinant IL-2 (2 ng/ml) (Gibco, USA), IL-7 (10 ng/ml) (Peprotech, USA) and IL-15 (1 ng/ml) (Peprotech, USA) were supplemented and repeat every two days. After two rounds of stimulation, neoantigens specific CTLs were harvested to carry out Cytotoxic assay and ELISPOT.

### (vi) ELISPOT assay

Enzyme-linked immunospot assay relies on quantifying IFN-γ-releasing to identify specific T cells. This method is similar to before(Depla *et al.*, 2008) and were performed using a Human IFN-gamma ELISpot PLUS kit (Mabtech, Sweden) according to the manufacturer’s instruction. Briefly,10ug/ml candidate peptides, 10ug/ml positive control peptides and 10ug/ml negative peptides were loaded on T2 or HCT116 cells respectively for 4h and co-incubate with specific CTLs at the ratio 1:4 (2500 T2 or HCT116 cells:10000 CTLs) in 96-well plate for 20h at 37°C, in humid 5% CO_2_ incubator. APOL1_176–184_ (ALADGVQKV) served as a negative control, Melan-A_26-35_ (ELAGIGILTV) served as a positive control. Next day, added the corresponding antibody and fluorescent developer to generate the spot after washing the plate 5 times. Countable spots were observed on the bottom plate represent the active cell that response to peptide-pulsed T2 or HCT116 cells. The standard we set are similar to previously: (number of experimental peptide spots)/(number of unrelated peptide spots)> 2 (Yamamiya *et al.*, 2018).

### (vii) Cytotoxicity assay

Cytotoxicity assay was performed with a CytoTox 96® Non-Radioactive Cytotoxicity Assay kit (Promega, USA) according to manufacturer’s instruction. We attempted to test the cytotoxicity of CTLs to target cells (T2 cells) loaded with 10ug/ml candidate peptides, negative peptides (APOL1_176–184_: ALADGVQKV) and without peptides. In our experiment, peptide-specific CTLs were co-cultured with T2 cells loaded with candidate peptides/negative peptides in different effector: target ratios (10:1 and 1:1) and incubated for 4h in 96-well V-bottom plate. We set control group according to instruction: effector cell spontaneous release group, target cell spontaneous release group, maximum release group, culture medium background control, volume correction control. After adding lysis solution(10x) for 45min in maximal release group and volume correction control, the plate was centrifuged at 250g for 4 minutes and transferred supernatant to a new plate. The substrate (contained in the CytoTox 96® kit) were added to each well, incubated for 30 min at room temperature and protected from light. Stop solution was used to terminate the reaction. Finally, we measured the absorbance at 490nm and recorded the value. Cytotoxicity percentage of effector cells = [(experimental release - effector cell spontaneous release - target cell spontaneous release)/ (maximum release - target cell spontaneous release)] *100%.

## 3. Result

### (i) HLA-A*0201-restricted neoantigens prediction in breast cancer

Our objective was to discern neoantigens derive from missense mutation base on neoantigen prediction system, which are capable of specifically activating CTLs to attack tumor and becoming new targets for immunotherapy of breast cancer in the future. We established a candidate epitopes prediction system similar to the method previously described (Johanns *et al.*, 2016). We identified 39,401 missense mutations from 969 patient’s tumor genomic data from TCGA, ICGC and an article. We determined 9,913 candidate epitopes (HLA-A*0201, length=9 mer) through the combination of six affinity prediction algorithms and the filter of IC50 < 500 nM in at least two algorithms(Fig.1*a*). The top 30 candidate epitopes were listed for further assessment at final base on mutation frequency and the mean of six binding score (Table 1). We found PIK3CA mutated in 8 of 40 breast cancers through our prediction system. As the previous description, high frequency of mutations of oncogenic PIK3CA gene are discovered in quite a few cancers, including lung cancer, breast cancer and colorectal cancers(Yardena *et al.*, 2004). This result supports the accuracy of our affinity prediction pipeline to a certain extent.

**Fig. 1.**
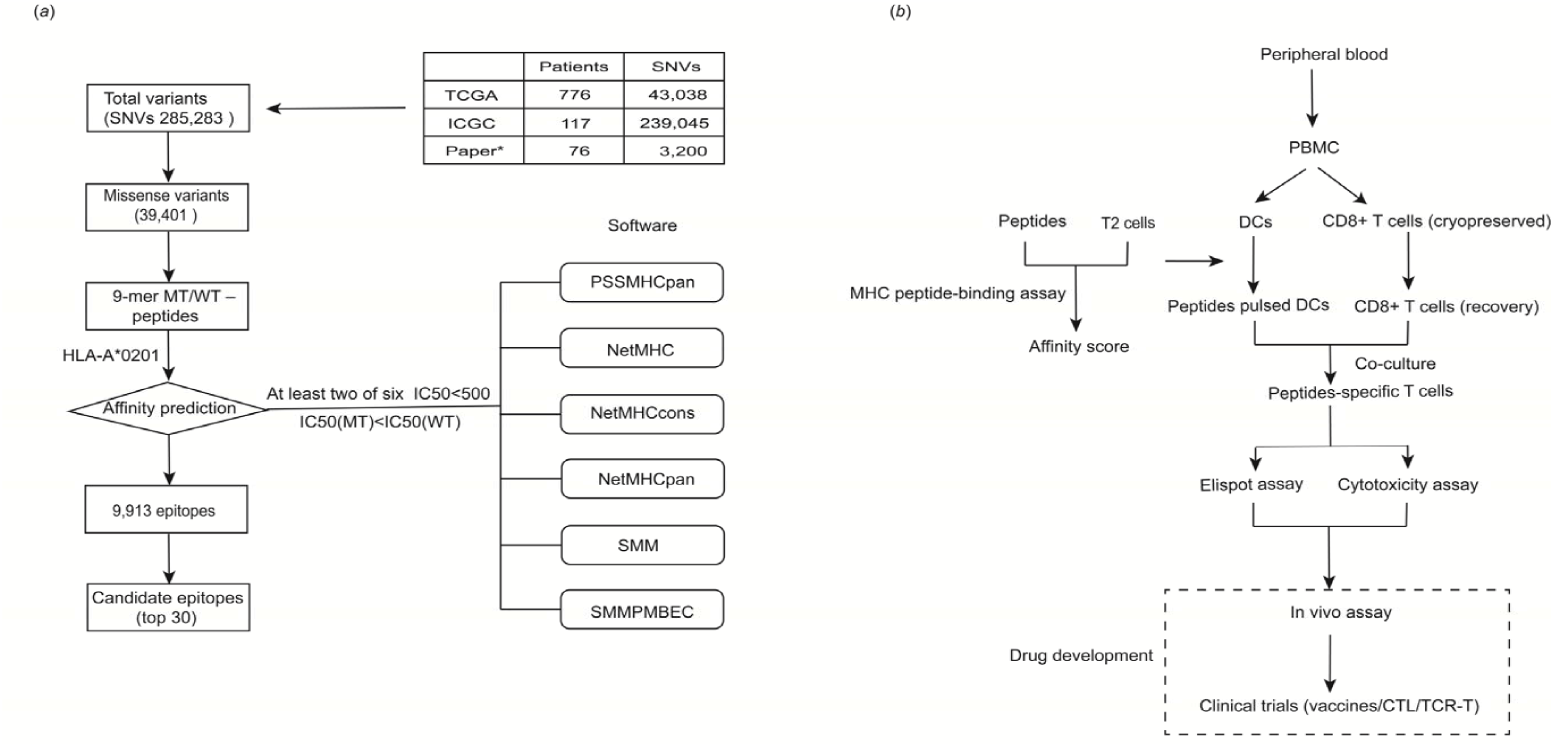
Flowchart of high-frequency neoantigen validation. (*a*) Pipeline of breast cancer high-frequency neoantigen bioinformatic identification. SNVs, single nucleotide variations; WT, wild type; MT, mutation type. (*b*) Neoantigen immunogenicity in vitro testing. Peripheral blood was from HLA-A*0201 healthy human donors. The content in the dotted box shows promising drug development. PBMC, the peripheral blood mononuclear cells.

**Table 1.** Top 30 epitopes from affinity predicted to bind HLA-A*0201

| Gene | Mutation position | Epitope | Score | NetMHC |  | NetMHCcons |  | NetMHCpan |  | PSSMHCpan |  | SMM |  | SMMPMBEC |  |
| --- | --- | --- | --- | --- | --- | --- | --- | --- | --- | --- | --- | --- | --- | --- | --- |
|  |  |  |  | MT | WT | MT | WT | MT | WT | MT | WT | MT | WT | MT | WT |
| **SF3B1** | p.K700E | GLVDEQQEV | 0.74 | 36.00 | 143.00 | 38.96 | 124.66 | 41.53 | 107.73 | 3.02 | 2.82 | 91.01 | 210.43 | 87.99 | 210.10 |
| **OPRK1** | p.R252H | LMILHLKSV | 0.75 | 42.00 | 114.00 | 34.40 | 116.20 | 28.14 | 118.49 | 2.94 | 2.36 | 71.63 | 122.21 | 76.11 | 131.65 |
| **KCNA1** | p.T226M | FIVEMLCII | 0.75 | 25.00 | 36.00 | 30.54 | 37.92 | 36.12 | 39.13 | 1.96 | 2.57 | 84.35 | 103.30 | 69.73 | 88.19 |
| **CBFB** | p.N104S | MILSGVCVI | 0.66 | 64.00 | 103.00 | 69.88 | 105.42 | 75.60 | 107.13 | 1.94 | 2.26 | 150.70 | 175.43 | 135.34 | 156.11 |
| **PIK3CA** | p.N345K | ILCATYVKV | 0.66 | 99.00 | 60.00 | 92.58 | 48.63 | 86.34 | 39.15 | 3.13 | 3.06 | 120.81 | 103.30 | 112.05 | 94.07 |
| **SPIK3CA** | p.E453K | GLKDLLNPI | 0.61 | 104.00 | 51.00 | 74.16 | 60.38 | 52.31 | 71.17 | 2.27 | 2.14 | 216.83 | 163.35 | 196.53 | 134.10 |
| **GPR32** | p.T332P | SLPSALARA | 0.46 | 190.00 | 167.00 | 172.47 | 160.76 | 156.62 | 154.48 | 2.44 | 2.30 | 299.99 | 340.49 | 323.17 | 352.72 |
| **PCDHA1** | p.T633M | GLYTGEISM | 0.45 | 160.00 | 693.00 | 173.40 | 476.88 | 187.63 | 327.56 | 1.71 | 1.64 | 316.31 | 634.03 | 315.09 | 634.50 |
| **PIK3CA** | p.G118D | ILNREIDFA | 0.23 | 650.00 | 1754.00 | 331.89 | 828.06 | 168.99 | 393.65 | 1.76 | 1.50 | 402.82 | 687.24 | 394.85 | 658.31 |
| **ERBB2** | p.V777L | MAGLGSPYV | 0.22 | 292.00 | 369.00 | 337.32 | 464.16 | 388.51 | 586.78 | 0.78 | 0.50 | 412.20 | 428.66 | 409.67 | 429.96 |
| **PIK3CA** | p.H1047L | FMKQMNDAL | 0.20 | 438.00 | 27837.00 | 328.32 | 25982.58 | 246.25 | 24223.28 | 1.35 | -1.64 | 444.74 | 31923.46 | 450.23 | 33762.14 |
| **ZNF766** | p.S147L | FISHSSSVL | 0.09 | 528.00 | 15637.00 | 466.68 | 15881.05 | 414.62 | 16116.46 | 1.43 | -1.14 | 422.78 | 7449.21 | 458.60 | 8287.62 |
| **RHO** | p.I263M | FLMCWVPYA | 0.83 | 1.00 | 2.00 | 1.70 | 2.06 | 1.46 | 1.82 | 6.12 | 5.58 | 0.62 | 1.59 | 0.49 | 1.27 |
| **CWC22** | p.M339I | YMIEVIFAV | 0.83 | 2.00 | 2.00 | 1.69 | 1.73 | 1.40 | 1.45 | 6.04 | 5.52 | 0.33 | 0.46 | 0.31 | 0.43 |
| **HVCN1** | p.K157N | FMMEIIFNL | 0.83 | 2.00 | 2.00 | 1.71 | 1.79 | 1.45 | 1.51 | 6.08 | 6.15 | 0.39 | 0.46 | 0.37 | 0.45 |
| **CCNA2** | p.V219F | ILVDWLFEV | 0.83 | 2.00 | 2.00 | 1.79 | 2.38 | 1.54 | 2.02 | 5.74 | 5.15 | 0.83 | 2.73 | 0.70 | 2.31 |
| **TMEM59L** | p.D315N | FMMEPNWPL | 0.83 | 2.00 | 2.00 | 1.65 | 1.81 | 1.35 | 1.49 | 5.36 | 4.84 | 1.00 | 1.61 | 0.99 | 1.65 |
| **MRGPRX4** | p.G233V | FLLCGLPFV | 0.83 | 2.00 | 46.00 | 1.96 | 36.31 | 1.73 | 28.26 | 6.33 | 2.61 | 0.74 | 52.25 | 0.66 | 46.93 |
| **KCNJ10** | p.V84M | FLFGMVWYL | 0.83 | 2.00 | 2.00 | 2.04 | 2.02 | 1.77 | 1.73 | 6.37 | 6.87 | 0.72 | 0.53 | 0.64 | 0.49 |
| **TMC5** | p.S744T | LLMDVFVTL | 0.83 | 2.00 | 2.00 | 2.14 | 2.10 | 1.92 | 1.91 | 6.69 | 6.28 | 1.09 | 0.95 | 0.99 | 0.86 |
| **OR5T1** | p.T50P | FLAIYLFPL | 0.83 | 2.00 | 4.00 | 2.59 | 4.26 | 2.52 | 3.94 | 5.18 | 5.86 | 0.86 | 1.63 | 0.84 | 1.65 |
| **TAS2R38** | p.P204T | YLWSVTPFL | 0.83 | 2.00 | 3.00 | 2.22 | 2.59 | 2.04 | 2.13 | 6.23 | 6.16 | 1.40 | 1.64 | 1.41 | 1.67 |
| **SLC13A3** | p.S377Y | FLYDAVTGV | 0.83 | 2.00 | 3.00 | 2.25 | 2.81 | 1.84 | 2.63 | 6.06 | 5.32 | 1.54 | 3.47 | 1.44 | 3.34 |
| **CMTM5** | p.C153F | FLIACAFLV | 0.83 | 2.00 | 8.00 | 2.83 | 11.53 | 2.98 | 15.05 | 4.64 | 3.41 | 0.79 | 6.81 | 0.75 | 6.72 |
| **CLN3** | p.H120Y | YLLPYSPRV | 0.83 | 2.00 | 4.00 | 2.08 | 4.62 | 1.84 | 4.60 | 5.90 | 4.64 | 1.80 | 15.46 | 1.83 | 15.05 |
| **KIDINS220** | p.R638L | FLATRLFLV | 0.83 | 2.00 | 3.00 | 2.51 | 2.97 | 2.28 | 2.84 | 5.23 | 5.08 | 1.46 | 2.62 | 1.52 | 2.80 |
| **CDC37L1** | p.H203Y | YLILWCFYL | 0.83 | 2.00 | 4.00 | 2.97 | 4.38 | 2.97 | 4.51 | 5.83 | 5.55 | 0.98 | 2.46 | 0.91 | 2.41 |
| **TPTE** | p.D140G | FLMGVLLRV | 0.83 | 3.00 | 2.00 | 2.75 | 2.08 | 2.38 | 1.77 | 5.81 | 6.17 | 0.99 | 0.46 | 1.01 | 0.44 |
| **TWISTNB** | p.N136Y | KLMGIVYKV | 0.83 | 2.00 | 5.00 | 2.44 | 4.52 | 2.11 | 4.06 | 5.66 | 5.21 | 1.81 | 8.36 | 1.83 | 8.62 |
| **COL14A1** | p.S1040F | FMVDGFWSI | 0.83 | 2.00 | 2.00 | 2.03 | 2.17 | 1.74 | 1.92 | 4.76 | 4.44 | 2.48 | 3.78 | 2.30 | 3.59 |

Note : score, the IC50 standardized as binding score. PSSMHCpan, a novel software based on PSSM.

### (ii) Binding of candidate peptides to HLA-A*0201

To further confirm the binding affinity of candidate peptides, peptides were synthesized and T2 cells peptide binding assay was test. T2 is a cell line deficient in transporter associated with antigen processing (TAP), causing endogenous antigens fail to present and HLA class I molecules on the cell surface to be unstable and prone to degradation. But the exogenous peptides are opposite, HLA class I molecules become stable. Therefore, T2 cells were used to detect the ability of candidate peptides binding to HLA. The higher the binding ability, the higher the expression level of MHC molecules on the cell surface(Hansen & Myers, 2003). We screened 10 high-affinity candidate peptides with FI>1.5 from 30 candidate peptides by T2 affinity detection to further perform ELISPOT assay and Cytotoxicity assay (table.2).

**Table 2.** The HLA-A2 binding affinity of candidate epitopes was confirmed by T2 binding assay

| Gene | Epitope | MFI (candidate peptide) |  | MFI (without peptide) |  | FI | Affinity |
| --- | --- | --- | --- | --- | --- | --- | --- |
| **RHO** | FLMCWVPYA | 1742.83 | 1750.01 | 439 | 439 | 2.98±0.008 | High |
| **SLC13A3** | FLYDAVTGV | 822.11 | 807.75 | 229 | 225 | 2.59±0.045 | High |
| **PIK3CA** | GLKDLLNPI | 880.68 | 859.2 | 246 | 240 | 2.58±0.063 | High |
| **CLN3** | YLLPYSPRV | 778.6 | 765 | 229 | 225 | 2.40±0.042 | High |
| **PIK3CA** | ILCATYVKV | 829.02 | 808.8 | 246 | 240 | 2.37±0.059 | High |
| **MRGPRX4** | FLLCGLPFV | 712.19 | 699.75 | 229 | 225 | 2.11±0.039 | High |
| **COL14A1** | FMVDGFWSI | 762.6 | 744 | 246 | 240 | 2.10±0.054 | High |
| **TWISTNB** | KLMGIVYKV | 693.72 | 676.8 | 246 | 240 | 1.82±0.049 | High |
| **CCNA2** | ILVDWLFEV | 590.82 | 580.5 | 229 | 225 | 1.58±0.032 | High |
| **CDC37L1** | YLILWCFYL | 583.95 | 573.75 | 229 | 225 | 1.55±0.032 | High |

Note : FI = (mean FITC fluorescence with a candidate peptide - mean FITC fluorescence without peptide) / (mean FITC fluorescence without peptide). When FI >1.0 indicates high-affinity peptides. FI values were shown as the mean ± standard deviation (n=2). FI, Fluorescence index.

### (iii) The identification of neoantigen immunogenicity in vitro

To identify the immunogenicity of these 10 neoantigens with high affinity, CD8^+^ T cells were separated from HLA-A*0201 healthy human donators and were stimulated in vitro by DCs loaded with these ten candidate peptides. The experiment process is shown in Fig.1*b*. After two rounds of stimulation, neoantigen specific T cells were amplified and used to perform IFN-γ ELISPOT assay and Cytotoxicity assay. Seven of ten (70%) candidate peptides induced specific CTLs response when compare the number of IFN-γ between candidate peptides and unrelated peptides (APOL1) (Figs. 2 *a* and *b*). Two peptides were derived from the PIK3CA among these 7 peptides. Among these 7 immunogenic peptides, TWISTNB-derived peptide (KLMGIVYKV) induced IFN-γ significantly increased compare to other candidate peptides, (number of experimental peptide spots)/(number of unrelated peptide spots)> 25.

**Fig. 2.**
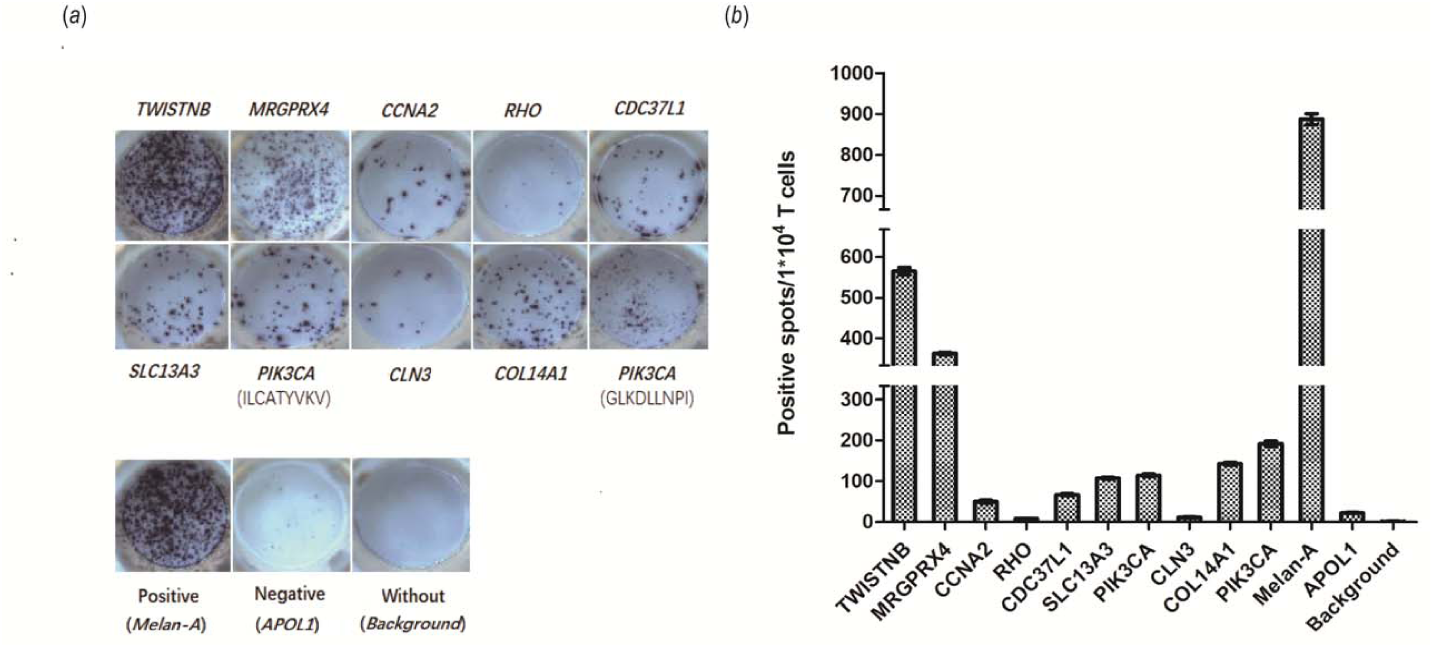
Peptides’ immunogenicity testing by ELISPOT assay. (*a*) Photographs from ELISPOT of specific-CTLs stimulated with T2–restricted candidate neoantigens. Images of ELISPOT wells show the IFN-γ released/10000 cells. Peptides are labeled with their corresponding gene name (shown in Table 1). (*b*) Each bar represents the mean ± SD from number of IFN-γ spots of two duplicate wells.

Furthermore, we tested the cytotoxicity of neoantigen specific T cells via Cytotoxicity assay. T2 cells loaded with neoantigens as the target cells were applied in cytotoxicity assays. There are 5 peptides (50%) can induce specific CTLs respond to the peptide specific pulsed T2, including *TWISTNB, PIK3CA, COL14A1, SLC13A3* and *CDC37L1* (Fig.3). As we expected, these specific CTLs could lyse the T2 cells pulsed with candidate peptides more efficiently than Irrelevant peptides, and these five peptides were concluded in those 7 specific peptides tested in ELISPOT. This result confirmed that these predicted neoantigens can cause specific T cell immune response and destroy target cells specifically in vitro. Similar to the results of ELISPOT, we observed the peptide KLMGIVYKV derive from TWISTNB has the strongest ability to induce T cells respond and lyse target cells (the percentage of specific lyse increase about 35% from 1:1 to 10:1) in those 5 immunogenic peptides.

**Fig. 3.**
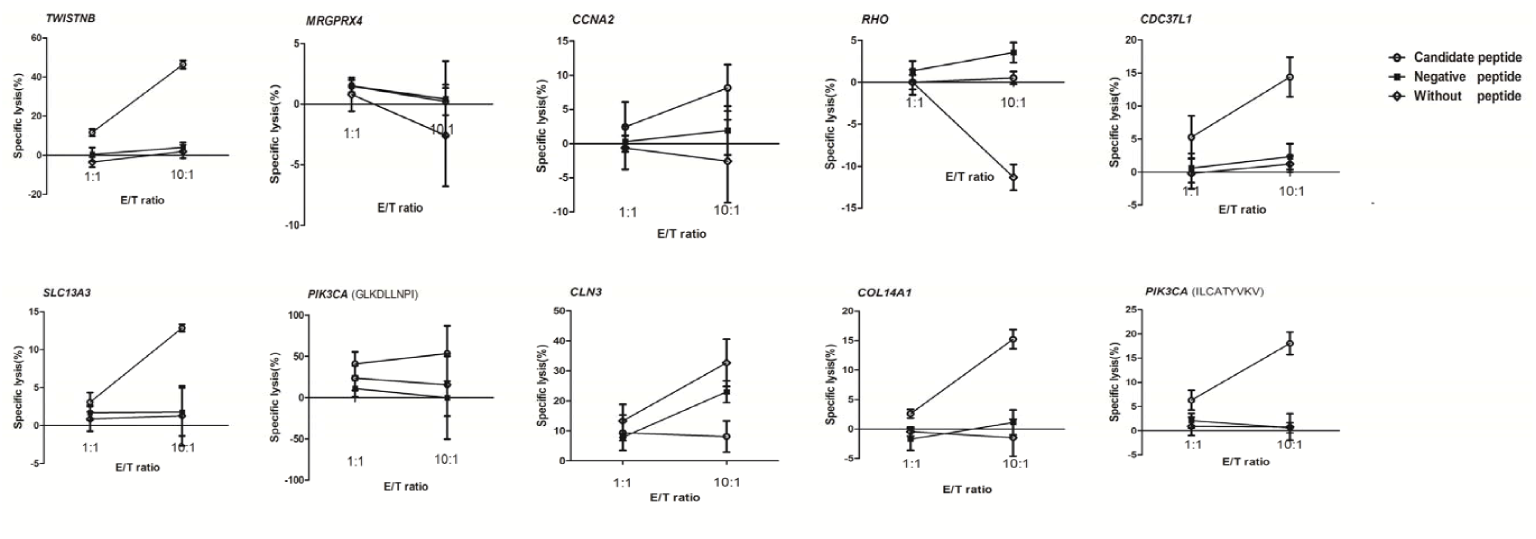
Specific-CTLs cytotoxicity assay measured with T2 cells. Peptides specific-CTLs cytotoxicity was detected at different effector/target ratios (1:1 and 10:1) against T2 cells pulsed with candidate peptides. Negative peptide, T2 pulsed with peptide(*APOL1*). Experiments were repeated three times and data represent mean±SD.

### (iv) Neoantigen from TWISTNB

To further describe the specificity of neoantigen from TWISTNB and confirm the cross-immune response in wild type (WT)-peptide and mutant type (MT)-peptide, we generated the MT-peptide/WT-peptide specific CTLs respectively in vitro and carried out ELISPOT assay. As the result, MT-peptide specific CTLs can recognized T2 cells loaded with MT-peptide and secret IFN-γ but not responded to the wild-type peptide pulsed T2 (p<0.01) (Fig.4*a*). Similarly, the WT-peptide specific CTLs can only respond to WT-peptide pulsed T2 in ELISPOT assay (Fig.4*b*). Above results indicated that the TWISTNB-derived neoantigen can be recognized by specific-CTLs and there is no cross-immune response with wild-type. We next used TWISTNB-induced CTLs to target genetically modified HCT116 cells and wild-type HCT116 cells in ELISPOT assay. Results shown that compared with wild-type HCT116 cells, specific CTLs secreted IFN-γ significantly increase (p<0.01) (Fig.4*c*), indicated that the neoantigen derive from TWISTNB can present on the surface of tumor cell line and identified by specific-CTLs. In further verification, we found that neoantigen from TWISTNB can induce immune responses in 7 random different donators in vitro(P<0.05) (Fig.5). We suppose that this neoantigen may serve as a positive control for the prediction of neoantigen in breast cancer and as the target for immunotherapy, such as DC-CTLs, neoantigen cancer vaccine and TCR-T (T cell receptor).

**Fig. 4.**
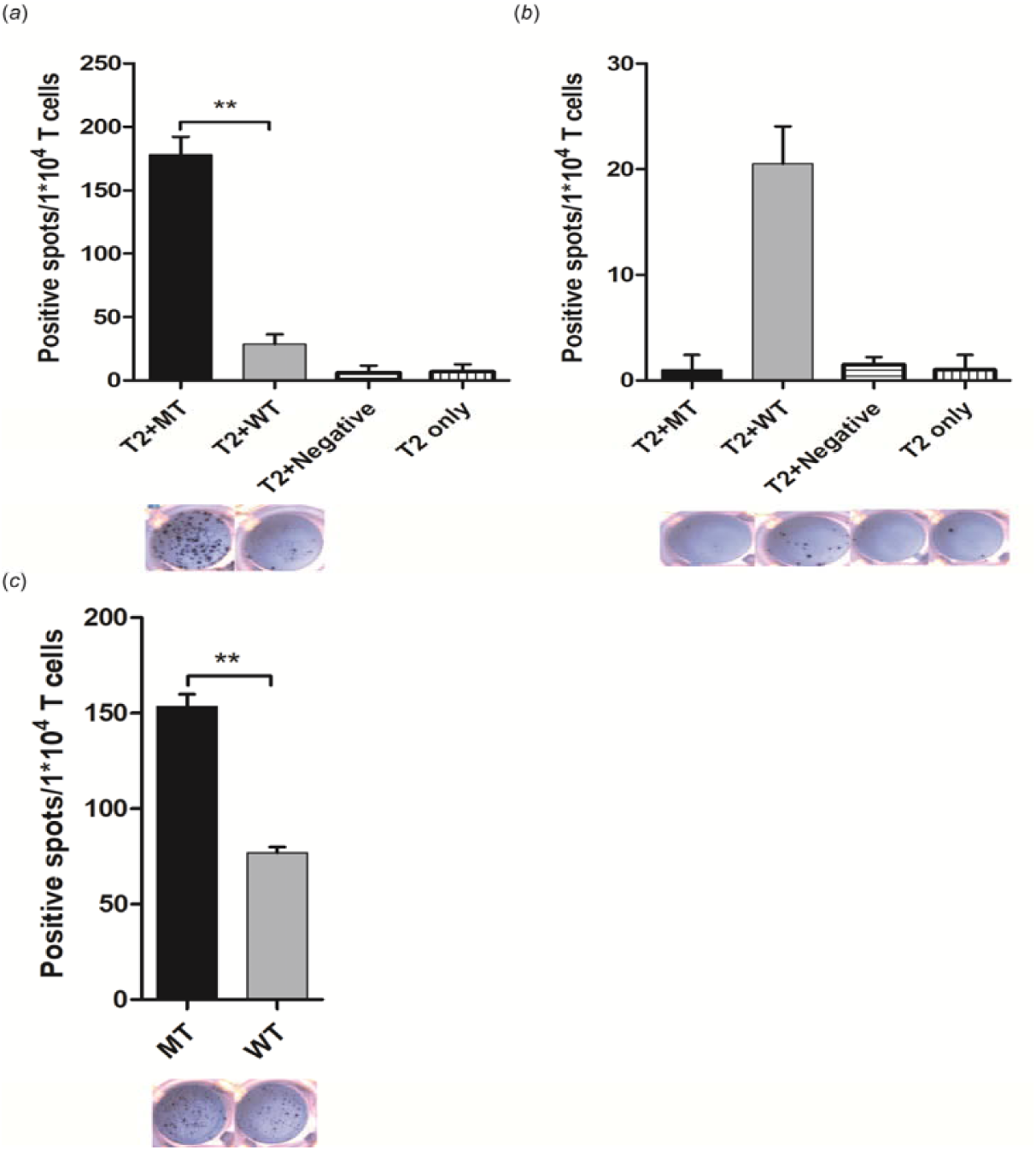
Peptide derive from TWISTNB was confirmed. (*a*) MT peptide (TWISTNB)-specific CTLs were stimulated by T2 with MT and WT peptide derived from TWISTNB respectively in the IFN-γ ELISPOT assay. (*b*) WT peptide (TWISTNB)-specific CTLs were stimulated by T2 cells with MT and WT peptide derived from TWISTNB respectively in the IFN-γ ELISPOT assay. (*c*) MT peptide (TWISTNB)-specific CTLs were stimulated by HCT116 with MT and without peptide derived from TWISTNB respectively in the IFN-γ ELISPOT assay. The data of histogram derived from the statistics of the spot number in ELISPOT well below. T2+Negative, T2 pulsed with irrelevant peptide(*APOL1*). One-tailed Student’s *t* test (\**P* < 0.05, \*\**P* < 0.01).

**Fig. 5.**
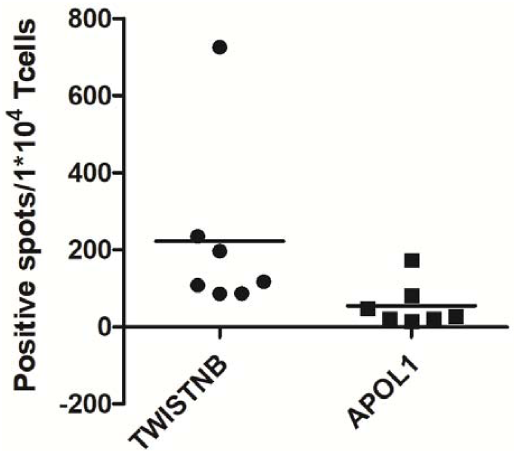
Immunogenicity verification of mutation peptide derived from TWISTNB in 7 healthy donors. Peptide specific-CTLs were generated from whole blood of 7 healthy donors and assayed by IFN-γ ELISPOT. Considering result as positive when the ratio of arithmetic means of candidate peptide to irrelevant peptide is ≥ 2.0. One-tailed Student’s *t* test (*P*=0.0425).

## 4. DISCUSSION

In our study, we obtained 285,283 SNVs contain 39,401 missense mutations from breast cancer samples published on authoritative databases and journal for predicting neoantigens base on 6 algorithms and further analyzed the immunogenicity of candidate neoantigens in vitro. We conducted cell experiments in vitro using T cells of healthy donors for filtering precise candidate neoantigens, 7/10 neoantigens were immunogenic consistent with ELISPOT assay criteria, 5/10 were capable of inducing specific T cells to cleave target cells loaded with neoantigens, these five immunogenic peptides were contained in those 7 neoantigens verified by ELISPOT. For mutation-type peptide derive from TWISTNB, ELISPOT and Cytotoxicity assay revealed stronger specific immune responses compare with other neoantigens, without cross-reactivity against the wild-type. In addition, it also demonstrated immunogenicity among 7 random health donors.

It is not difficulty to detect genome in the era of rapid development of sequencing and information technology. The genomic information of tumors can be obtained from tumor clinical samples or databases which record various cancer information. In terms of individualized vaccines base on neoantigen, which tailored to patient tumors, are able to target heterogeneity tumor cells and trigger strong Immune response(Hellmann & Snyder, 2017) or (Chu *et al.*, 2018). Although the efficacy is remarkable, neoantigens were predicted are individualized and cannot be popularized when encountering multiple cancers and multiple patients in clinical practice. On various database platforms, such as TCGA, ICGC, etc., the normal and mutant information of tumors can be found, providing a great resource for studying evolution and progression in tumors. Some studies indicate that based on the cancer database, we can research the relationship between the mutation type and development of tumors, predict neoantigen and establish neoantigen database, develop software and other events(Wu *et al.*, 2018) or (Turajlic *et al.*, 2017) or (Zhou *et al.*, 2017). For some researchers, obtaining tumor-related information from database is reliable, efficient, and time-saving for confirming neoantigens. So, we selected the method to find neoantigens. In spite of mining database information makes a huge contribution to predicting neoantigens, it is necessary to perform experiments in vivo and in vitro for verifying neoantigens that are beneficial to tumor vaccines, TCR-T, DC-CTLs and other immunotherapies. Hence, we optimized the candidate neoantigens by cell experiments in vitro based on this consideration.

We did not choose the patient’s peripheral blood or tumor cells to screen neoantigens, but the peripheral blood of healthy donors. Strønen et al., in 2016, demonstrated that using T lymphocytes from healthy donors to validate tumor-specific neoantigens, establishing T cells repositories that can specifically identify tumor neoantigens, and providing more possibilities for discovering new targets for tumors immunotherapy(Strønen *et al.*, 2016). Other researchers also used T cells from healthy donors in the study of tumor-specific T cell receptors for avoiding such an event that the existing T cells are prevented from being activated again or initiating specific T cells in patients(Kato *et al.*, 2018). It can be seen that it is effective to use healthy donor T cells to optimize neoantigens. In our study, we identified a neoantigen derive from TWISTNB in breast cancer using healthy donator T cells and the neoantigen maybe applied in TCR-T, DC vaccine, peptide vaccine and DC-CTLs and others anti-tumor therapy.

Tumor formation is a process caused by normal cell dysfunction and genetic variation accumulate. These events lead to the expression of surface antigens such as tumor neoantigens and differentiation antigens(Chen & Mellman, 2013). The process of tumor immune response begins with antigen presenting cells recognizing tumor antigens, and then processes these antigens which will be present to T cells that arousing the immune response. Tumor antigens give rise to extensive attention at domestic and foreign, as one of the significant factors in anti-tumor immunity. There are two kinds of tumor antigens that are spontaneously recognized by immune cells: one is tumor-associated antigen, and the other is tumor mutant antigen(Coulie *et al.*, 2014). Over the last few decades, nonmutated tumor antigens vaccines have been extensively studied, but the clinical effects were not satisfactory. Rosenberg et al. summarized the clinical effects of various peptide vaccines in 2004 such as MART-1, gp100, NY-ESO-1 or Her2/neu, the overall objective response rate of clinical trials was only 2.9%(Rosenberg & Yang, 2004). The neoantigen, as a marker for identifying tumors, has been previously shown to be able to induce MHC restricted T cell responses in human(WöLfel *et al.*, 1995). In addition, neoantigens not only is important target of check point blockade therapy (Gubin *et al.*, 2014), but have an effect on the development of cancer vaccines which will play a role in clinical practice. Tran et al. have pointed out that neoantigen associated with KRAS mutations have been identified to induce tumor infiltrating lymphocytes in patients with colorectal cancer to subside tumors, indicating the enormous potential of neoantigens vaccines in tumor therapy (Tran *et al.*, 2016). Recently, when faced with the problem of heterogeneity in tumor immunotherapy, scientists began to focus on the research of individualized vaccines based on neoantigens, have been used for clinical. More inspiringly, a phase I clinical trial of personalized neoantigen vaccine for patients with melanoma was announced at Nature in 2017 (Ott *et al.*, 2017). Six patients with melanoma resection were vaccinated, and four patients had no recurrence during the follow-up period (20-32 months). More than that, a similar research result was made public in the same year(Sahin *et al.*, 2017). Anyhow, neoantigen is a very promising biomarker for tumor immunotherapy, and it is urgent to develop. In fact, only a small subset specific neoantigens can trigger immune response (Zhang *et al.*, 2017). In this study, a large number of breast cancer sequencing data published on authoritative databases and magazines were used for bioinformatics analysis and cell experiments. And we discovered a neoantigen derive from TWISTNB, which can induce immune response in 7 random individuals. With this neoantigen, we will devote to the development of tumor vaccines, TCR-T, DC-CTLs and other tumor immune cell therapies. With this pipeline, we can strive to achieve an objective that find shared biomarker that can target recurrent, metastatic and refractory breast cancer. It is beneficial to explore a high-frequency neoantigen target that can be shared in multiple cancers, identify abundant high-frequency neoantigens and create neoantigen library for immunotherapy.

## 5. Acknowledgement

This study was supported financially with funds from Science, Technology and Innovation Commission of Shenzhen Municipality (JCYJ20170303151334808), National Natural Science Foundation of China (81702826), Shenzhen Municipal Government of China (20170731162715261) and Yunnan Provincial Science and Technology Department (Le Cheng, grant Nos. 2014HB053 and 2016RA037).

